# Microbial single-cell omics in situ

**DOI:** 10.1101/2024.07.07.602369

**Authors:** Xihong Lan, Qiaoxing Liang, Jinhua He, Jiayi Wu, Xiaoying Zhang, Fei Li, Guoping Zhao, Ruidong Guo, Lili Li, Huijue Jia

## Abstract

Metagenomics has enabled understanding of the microbial composition and functional potential in various environments. Using Laser-Induced Forward Transfer (LIFT) technology, we report high-quality microbial single-cell genome or transcriptome in complex samples such as saliva, mouse gut and human tumor sections. Neighboring cells could be separately retrieved and bacteria could be fluorescently labelled. We illustrate cell-fate commitment with sporulation. The method is scalable and precise, and would empower fundamental insight for microbial populations and single-cell interactions with the host.

## Introduction

Single-cell transcriptome analyses for mammalian cells are reaching their prime, further boosted by spatial omics technology^1^. Single-cell omics for bacterial cells, however, have remained a daunting task, due to the orders of magnitude lower concentration of nucleic acids, the lack of a poly(A) tail, and the Poisson distribution of droplet microfluidics ^2^. Marine cyanobacteria were among the first bacteria to be analyzed using single-cell genomes ^3^, due to their dominance in the global ocean. Single-cell genomes of extreme environments have revealed genomic features of uncultured bacteria and archaea *^4^*. Other than low-diversity communities, microbial single-cell studies have mostly been illustrated for model organisms such as *E. coli*, and the genome or transcriptome analyzed, often using targeted methods, has remained far from complete *^5-9^*.

Laser-Induced Forward Transfer (LIFT) is a straight-forward way of manipulating microparticles including cells *^10-13^*. With a laser beam that evaporates or deforms a thin metal coating^14, 15^, LIFT is ideally suited for picking particles in the size range of bacteria. It can potentially tolerate food particles, cell debris, and other objects commonly present in the human microbiome that could undermine analyses using droplet microfluidics.

Metagenomic analyses of low-biomass environment such as tumors and placenta has been costly and controversial *^16^*. Thus, a direct approach to characterize the microbes in tissues with over 99.9% host sequences would also be very much welcomed.

Here we present microbial single-cell genomes and transcriptomes using a commercially available optics equipment, and demonstrate their superb utility in a number of scenarios.

## Results

### Draft genome quality assembly for microbial single cells

Consisted of several key modules^17^, our LIFT equipment was capable of image capture, Raman spectral acquisition and single cell isolation from complex samples after previous processing steps (Sup. Fig. 1a-d; Sup Movies S1). Next, we explored the energy and mode required for microbial cell ejection using bacteria of different sizes and shapes. Low energy resulted in cell ejection failure including cell shifting and cell rupture (Sup. Fig. 2a and b), while appropriate energy for cell ejection successfully isolated the selected cell without affecting the closely adjacent cells (Sup. Fig. 2c). Further separation of cells with different morphological characters suggested that appropriate energy for cell LIFT was related to the size of the cell rather than its shape (Sup. Fig. 2d). For the larger bacteria, multiple LIFT attempts could also successfully separate the cells without increasing energy (Sup. Fig. 2e).

To develop microbial single-cell technology for complex samples, we started with saliva samples which are even more diverse than feces. A fresh saliva sample was centrifuged and spotted onto the slide (Sup. Fig. 1c), and bacteria cells were visualized before LIFT (Fig1.a). Single cells were ejected by LIFT without affecting neighboring cells (Fig1.a and b, no burnt circle with the low energy 60nJ) and Multiple Displacement Amplification (MDA) was performed in a PCR tube. Without the need for sophisticated algorithms to pool assembly from fragmental genomes, as was reported recently *^5^*, here we used state-of-the-art algorithms developed for metagenomes *^18^*, and obtained a bacterial genome of 99.99% completeness and low contamination 0.33% (Fig1.c, Sup. Fig. 3a-d). This genome was identified as *Megamonas funiformis*. Gene annotation through Bakta revealed the detection of coding sequences (CDSs), tRNAs, rRNAs, ncRNAs, and CRISPRs (Fig1.d, Table S1). Calculating the coding density showed that the genome sequence is >90% coding (Sup. Fig. 3e and f), which is within the range for commensal bacteria (on average 87% for human gut *^19^*). Thus, LIFT from complex samples could obtain high-quality single-cell bacteria genomes (single-cell assembled genomes, SAGs) that far exceeds typical single-cell genomes ^5^ or metagenomically assembled genomes *^20^*, and are on-par with genomes from cultured isolates.

**Figure. 1.**
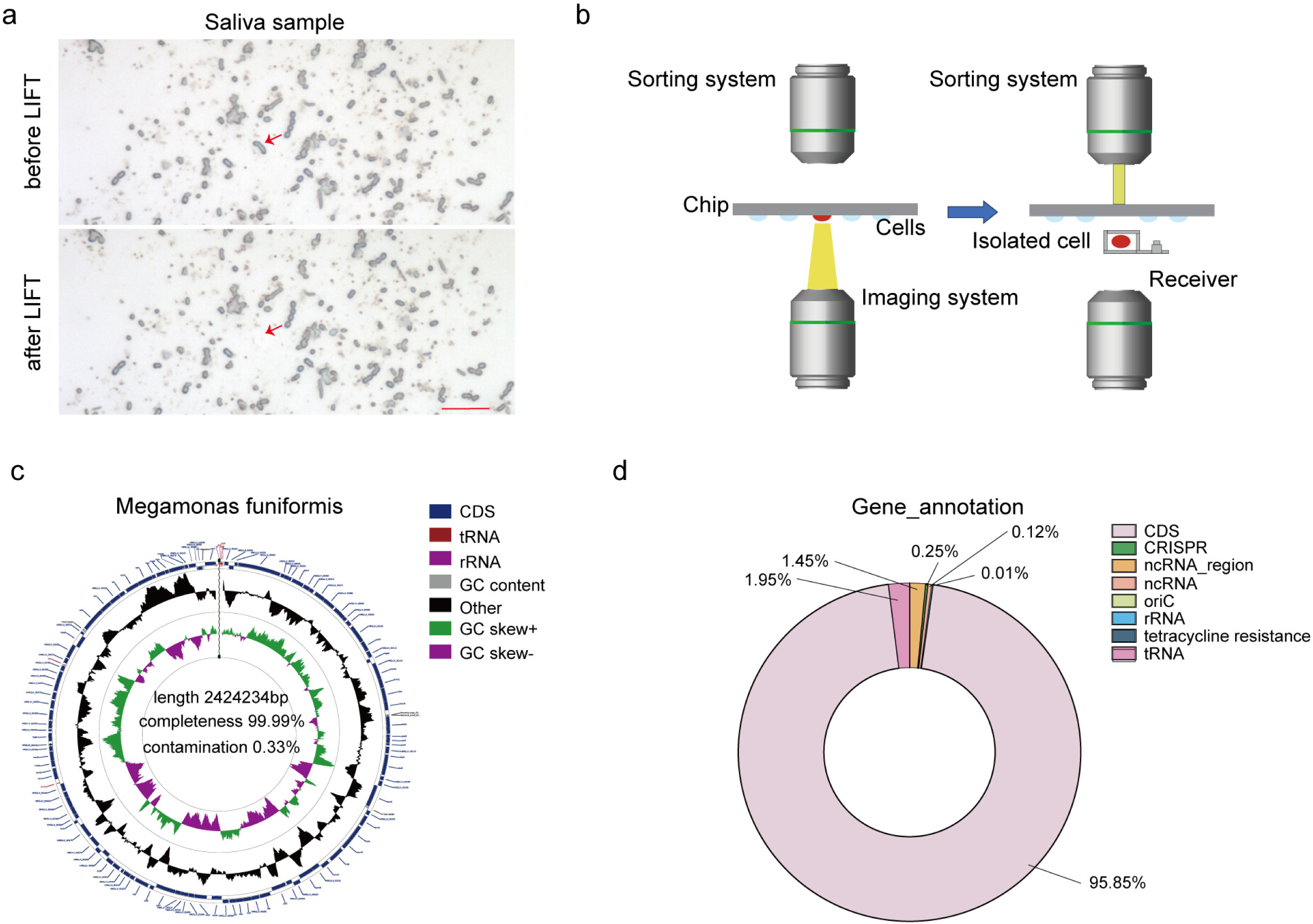
High-quality microbial single-cell draft genome obtained from saliva. **(a)** A view of the sample before and after ejection of the selected microbial cell. A single cell as indicated was selected for genomic sequencing. Scale bar 10μm. **(b)** Diagram illustrating the imaging by the inverted microscope and LIFT by the upright microscope, on the commercially available PRECI-SCS equipment. The collectors are placed on a wheel, whose rotation is controlled by the manufacturer’s software (Multiple cells could also be transferred into the same collector, depending on the experimental design). **(c)** Visualization of single-cell assembled genome by CGView tool displaying GC skew, feature labels, GC content, divider rings, and tick density. **(d)** The pie chart of single-cell assembled genome annotation display different categories including coding sequence (CDS), CRISPR, ncRNA region, ncRNA, replication origin (oriC), rRNA, tetracycline resistance, tRNA with their respective proportions.

To validate the automatic process, *Sphingomonas sp000797515* labeled with TADA were precisely ejected from sparse regions and densely packed regions in mixture of several species, sequenced using two different reaction volume, both of which yielded high-quality genomes (Sup. Fig. 4a-d; Sup Movies S2). There were no significant differences in genome completeness, contamination and coverage between the genomes obtained by the two methods suggesting that downsizing the reaction volume does not lead to a decline in SAGs quality (Sup. Fig. 4e). The single-cell genomes from LIFT were also highly consistent (ANI>95%) with the known input genomes (Sup. Fig. 4f), validating this approach.

To increase the throughput and eliminate individual differences in operation, we made use of an automated pipetting platform to take over the molecular biology work after automated single-cell ejection. With the aid of automatic identification by adjusting parameters such as diameter, area and fluorescence intensity, 34-62 bacteria from single strain suspensions and complex samples could be precisely ejected within one minute for the next step of library construction (Sup. Fig. 5a-c; Sup Movies S3-S4). Salivary microbes of three fields from a single donor were marked for collection (0.1s per cell), which consisted of diverse morphological characters (Fig2.a). After de novo assembly, 4 of the 39 SAGs have completeness >90 and contamination <5, which are regarded as high quality *^21^*. 27 of the 39 SAGs have completeness >50 and contamination <10, which are medium quality (Table S2). The species were identified according to GTDB-Tk with ANI >95%. With this relatively small-scale attempt, we already obtained oral bacteria from all major phyla, including *Actinobacteriota*, *Bacteroidota*, *Campylobacterota*, *Firmicutes*, *Fusobacteriota*, *Patescibacteria* and *Proteobacteria* (Fig.2b). Species from the oral cavity and related body sites were seen, such as *TM7x*, *Haemophilus sp.*, *Neisserria sp.*, *Porphyromonas gingivalis*, *Filifactor alocis*, *Fusobacterium peridonticum*, *Bifidobacterium vaginale* (renamed from *Gardnerella vaginalis*) and *Parabacteroides merdae* (Fig.2b). These results indicate that the single-cell genomes could be reliably obtained with medium to high quality in an automated and selective manner to reveal the microbial landscape in a complex sample.

**Figure. 2.**
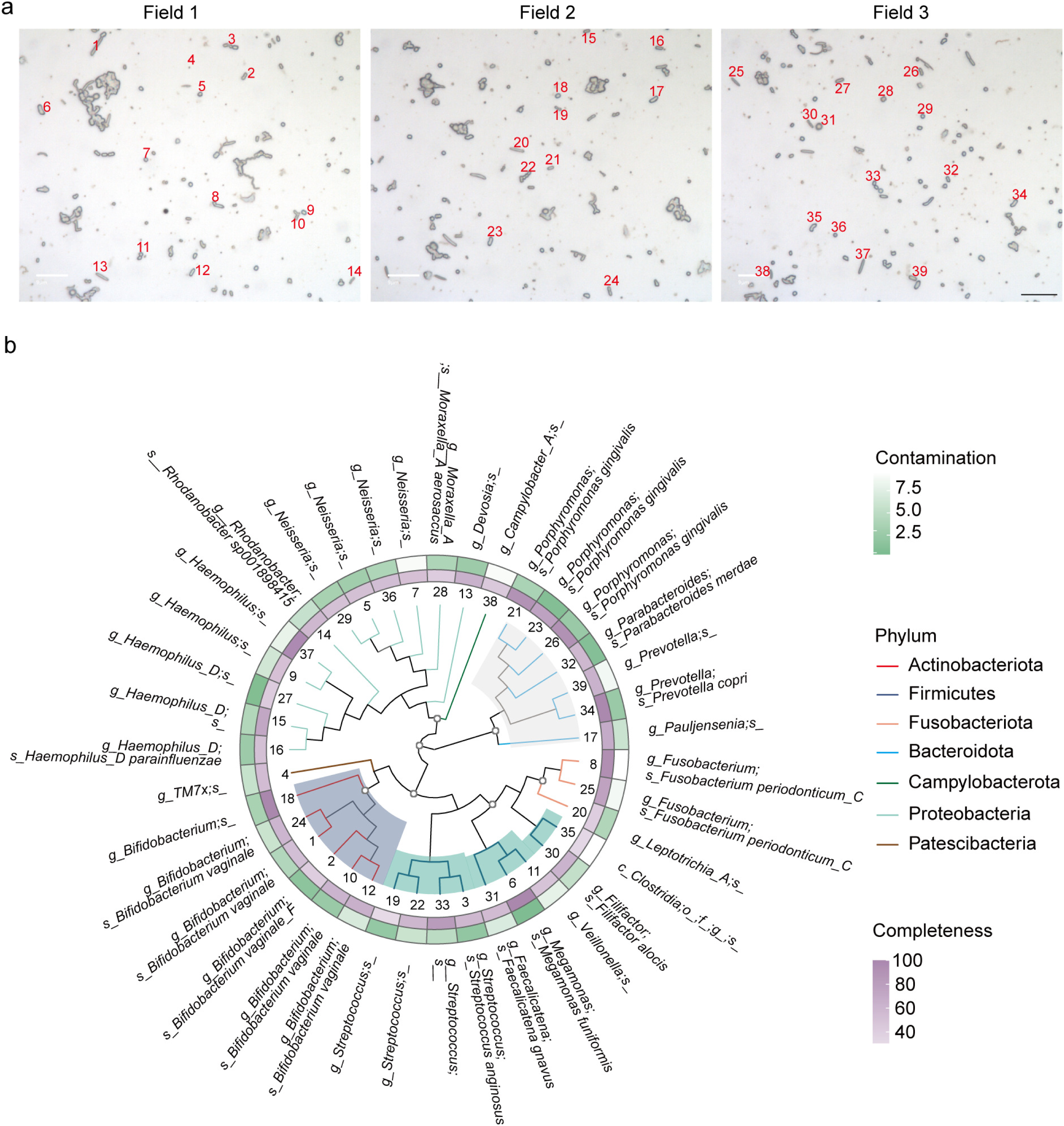
Automated microbial single-cell genomics. **(a)** Individual microbial cells of saliva distributed on the chip were observed before LIFT. The selected microbes numerically marked with variable morphology are ejected for single cell sequencing. Scale bar, 10μm. **(b)** A phylogeny constructed from 39 obtained SAGs genome sequence is represented by the inner dendrogram. The phylum of each SAGs is indicated by the branch color. Numbers for the tip nodes of the dendrogram are corresponding to numbers labeled in panel. Completeness and contamination of each SAGs are represented by two heatmaps overlay on the phylogenetic tree. The species name of each SAGs is listed in the outermost ring.

### Cell-fate commitment revealed by single-cell transcriptome

Sporulation is a strategy for bacteria to survive harsh environments ^22, 23^, and to potentially distribute to other hosts. We cultured a strain of *Bacillus licheniformis* isolated from human saliva, and LIFT-picked single cells to analyze their transcriptome following citric acid treatment (Sup. Fig. 6a). Thousands of transcripts were detected from cells stimulated by citric acid (Sup. Fig. 6b), which is consistent with widespread yet incomplete expression of the bacterial genome per cell cycle, known for model organisms such as *E. coli ^24^*. Besides protein coding genes, tRNAs and regulatory sRNAs were also detected (Sup. Fig. 6c), favoring the robustness of the experimental protocol.

In addition to scRNA analysis according to the bacteria reference genome (Sup. Fig. 6d-g), the high-quality reads obtained could also be de novo assembled. Gene counts obtained by these two analytical methods were similar (significance *p < 0.05*) and the expression profile was highly correlated (Sup. Fig. 6h). Thus, both known and unknown bacteria could be investigated using this scRNA approach.

Oligo(A) tails target bacterial mRNAs for degradation *^25^*; this coupling to exonucleases in bacteria is evolutionarily similar to exosome degradation of transcripts in the eukaryotic nucleus. We detected stretches of three or more adenylates in a number of transcripts (Sup. Fig. 7a-f, Table S3). TctA (Tripartite tricarboxylate transporter TctA family), a large transmembrane protein involved in citrate uptake, for example, showed long stretches of poly(A) spanning the entire gene, which led to three contigs in de novo assembly (Sup. Fig.7g and i). The shorter form was more abundant in five of the five cells analyzed (Sup. Fig.7h), consistent with degradation from 3’ to 5’. These results highlight the fundamental questions that can be addressed using high-quality microbial single-cell transcriptome.

We next explored the dynamic transcription changes during the commitment to sporulation in *B. licheniformis* following citric acid treatment. In addition to the upright laser for LIFT, the commercially available equipment we used also harbor an inverted laser for single-cell Raman spectrometry (Fig.3a). Both the cell morphology and Raman spectra could be rapidly obtained from a sample. During the citric acid-induced sporulation, the day 0, day 3 and day 9, *B. licheniformis* cells can be visually and spectrally divided into three groups, from day 0 rod (I), day 3 intermediate state (II) to day 9 oval spore (III) (Fig.3a and b). The mean spectrum of cells I, II, III were different, and spectra from the individual microbial cells formed three clusters (Fig.3c). Thus, the morphological features of bacteria as well as Raman spectrum indicated alterations in cellular gene expressions. Consistent with the decrease in growth, while ribosomal RNAs, ribosomal proteins, and nucleoid associated proteins dominated the day 0 transcriptome, the reads proportion of other genes notably increased as the cells undergo morphological changes towards sporulation (Sup. Fig. 8a). Comparison of the gene expression of cells I, II, III allowed us to identify 194 differentially expressed genes, with these transcripts mainly clustered into three groups according to variations in their reads counts (Fig.3d). Gene ontology (GO) analysis of Biological Process (BP), Molecular Function (MF), Cell Component (CC) showed that the genes preferentially expressed in the intermediate state of group II are linked to the pathways such as cellular response to stress, sporulation, hydrolase activity and intrinsic component of membrane (Sup. Fig. 8b). Consistently, electron microscopy confirmed that cellular component assembly occurred in cells II, which was also observed before Raman microscopy and LIFT (Sup. Fig. 8c). Comparing cells I and cells II, we found that germination gene *gerD* co-present with sporulation related genes such as spore coat protein and *YjcZ* family sporulation protein were upregulated after citric acid treatment at day 3 (Fig.3e). When cells I transformed into spore, *spoIIIAC* (stage III sporulation protein AC), *YtvI* (sporulation integral membrane protein) and spore-specific gene *splB* (spore photoproduct lyase) were highly expressed (Fig.3f).

**Figure. 3.**
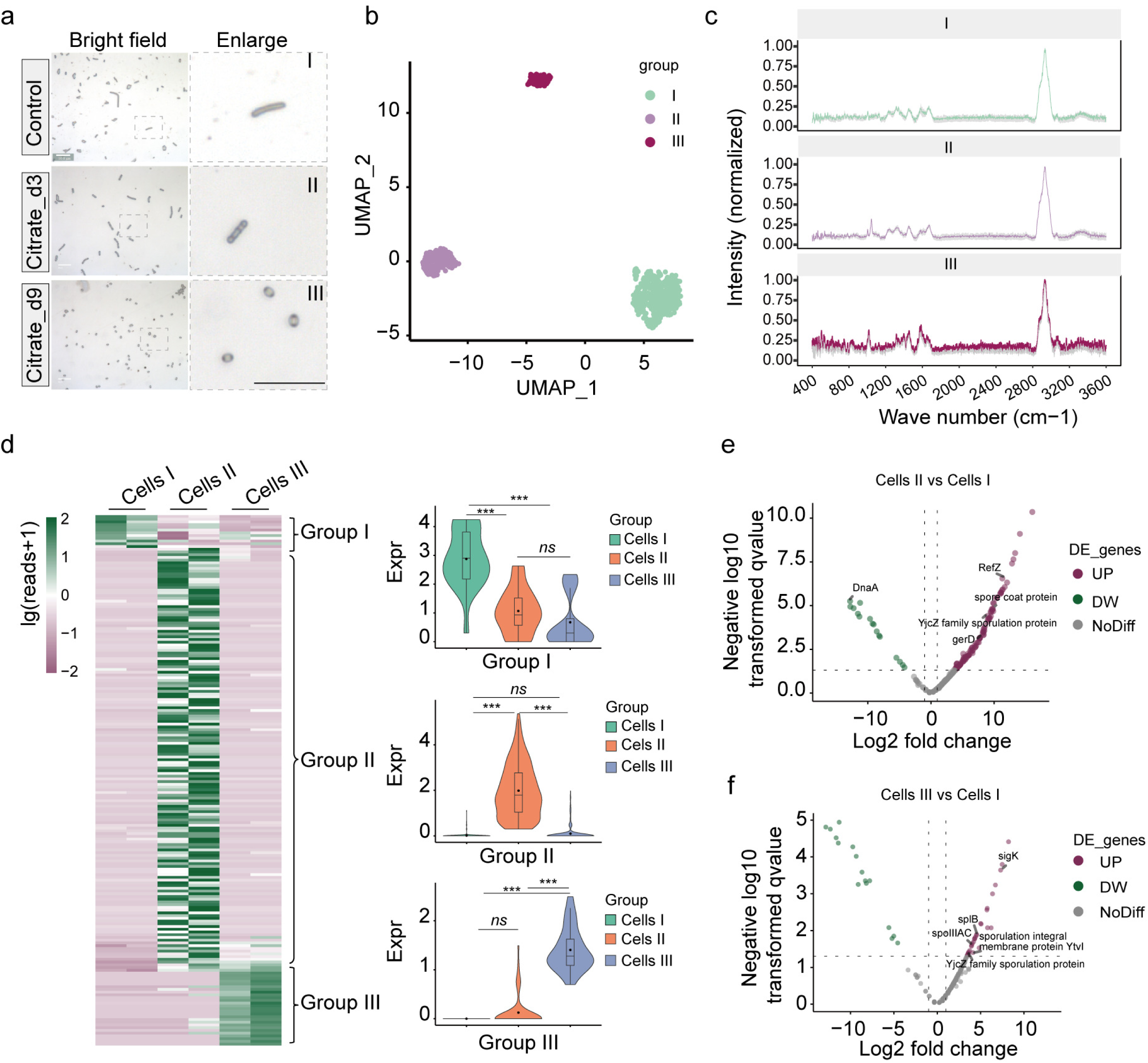
Coherent grouping of *B. licheniformis* cells visually, spectrally and with scRNA transcriptome. **(a)** Individual *B. licheniformis* cells I (treated without citric acid), II (treated with citric acid for 3 days), III (treated with citric acid for 9 days) distributed on the chip were observed in bright field before Raman spectrometry and LIFT. Scale bar, 10μm. **(b)** More than 50 cells from each group (Cells I, Cells II and Cells III) are selected for Raman spectra acquisition. Dimensionality reduction is performed on the obtained Raman spectra by UMAP. **(c)** The means and standard deviations of Raman spectra were calculated for each group. **(d)** *B. licheniformis* of morphological forms I (treated without citric acid), II (treated with citric acid for 3 days) and III (treated with citric acid for 3 days) were ejected (two cells for each group) for scRNA sequencing. Heatmap of differentially expressed transcripts in Cells I, Cells II and Cells III. The candidate transcripts are mainly clustered into three groups (left panel). Violin plot of normalized transcripts expression in Group I, Group II and Group III (right panel). Data are represented as mean ± SE (ns: *P* >0.05, *\*P* < 0.05, *\*\*P* < 0.01, *\*\*\*P* < 0.001) and calculated using ANOVA with post-hoc Tukey HSD Test. **(e)** Volcano plot of differentially expressed genes of Cells II versus Cells I. Gene counts normalized and compared by DeSeq2. **(f)** Volcano plot of differentially expressed genes of Cells III versus Cells I. Gene counts normalized and compared by DeSeq2.

Further looking into the day 3 samples led to five types of cells with distinct cell morphology, and the scRNA also formed five groups (Fig.4a). Principal Component Analysis of gene expression profiles revealed that cells of type I, type IV and type V were relatively clustered independently, while cells of type II and type III were relatively dispersed (Fig.4b). Comparing gene counts in each type, we found that cells of type II in cell division state were active in gene transcription (Fig.4c). Genes whose expression were enriched in the different cell types included enzymes, transcription regulator and biosynthesis proteins (Fig.4d). Genes highly expressed in type I are FdhF/YdeP family oxidoreductase, manA (mannose-6-phosphate isomerase, class I), PTS mannose/fructose/sorbose transporter subunit IIC, phosphoribosylaminoimidazolesuccinocarboxamide synthase, *ComGE* and 2-dehydropantoate 2-reductase, which enable type I cells resistance to low PH caused by citric acid, transporting sugar molecules, pantothenate biosynthesis, DNA uptake and purine biosynthesis ^26-30^. Genes highly expressed in type II cells included DEAD/DEAH box helicase, primosomal protein DnaI, DNA polymerase/3’-5’ exonuclease PolX, DNA helicase RecQ, RNA-binding protein and sigma-54-dependent Fis family transcriptional regulator, which indicated DNA replication and gene transcription process in cell division ^31, 32^. Genes observed in type II were also expressed in type III, but relatively lower. When cells changed into type IV, GTP associated genes MnmE, elongation factor Tu and GTP 3’,8-cyclase MoaA, 50S proteins and GntR family transcriptional regulator were highly expressed, which regulated biological process such as gene translation and cell motility ^33, 34^. Genes in type V were mainly transporters, Na+/H+ antiporter NhaC, permease and bacteriocin, which functioned in stress resistance, substance exchange and inhibition of spore germination ^35-37^. Comparing mean spectrum of five type cells, we observed Raman signal at band 750nm-785nm contributed by Cytochrome c and metabolites of purine degradation decreased (Fig.4e) ^38, 39^, which is consistent with variation of metabolic and gene transcriptional activity in five type cells. In addition, transcription of sporulation and spore related genes also varied with cell state (Fig.4f). These results illustrate the discovery power of high-quality scRNA, and indicate that visual and/or label-free Raman spectra-guided selection of microbial single cells could greatly facilitate the investigation of heterogeneous microbial populations. All cells could be rapidly scanned in seconds, before more costly amplification and analyses.

**Figure. 4.**
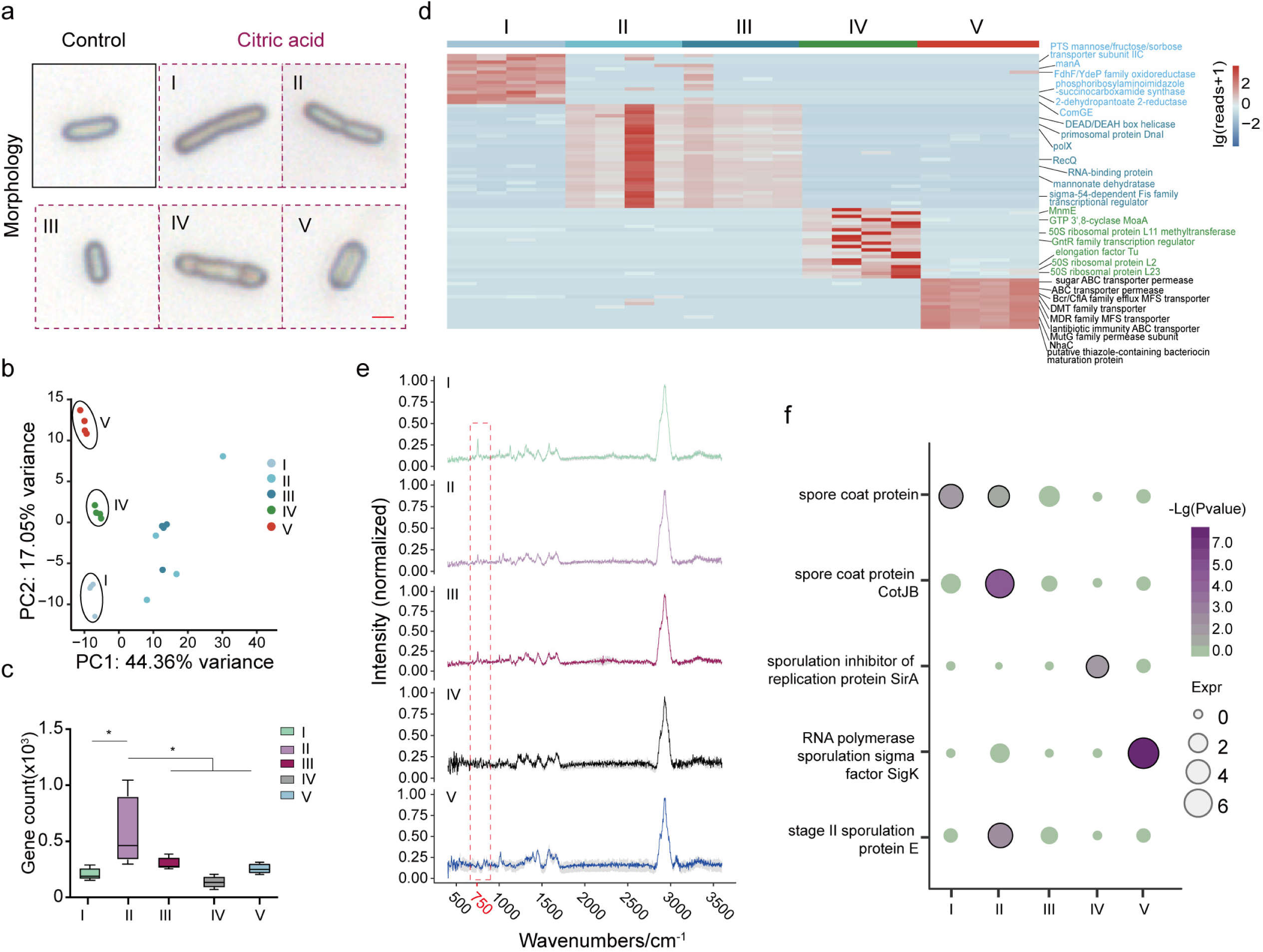
Single-cell transcriptome reveal heterogeneity in sporulation. (**a**) Image of five type (four cells for each type) *B. licheniformis* cells in sample treated with citric acid for 3 days and cells without citric acid treatment as control. Scale bar, 1μm. (**b**) Principal Component Analysis of gene expression profile from the five cell types. **(c)** Calculation of gene counts in each cell type. Data are represented as mean ± SE (*\*P* < 0.05, *\*\*P* < 0.01, *\*\*\*P* < 0.001) and calculated using two-tailed unpaired Student’s t-test. **(d)** Heatmap of representative gene expression (z-score of log-transformed values) from 20 individual *B. licheniformis* cells grown in day 3 citric acid-treated media. (**e**) Raman spectra of five type *B. licheniformis* cells. The spectra are averages of ten individual spectra and normalized to the intensity of the strongest feature in each spectrum. **(f)** Dot plot of genes involved in *B. licheniformis* sporulation. Gene expression and pvalue were calculated between the selected cell type and the remaining cell type.

### Single-cell microbial genome and transcriptome from tissue-resident **microbiota *in situ***

Having demonstrated single-cell draft genomes from complex samples and high-quality precision single-cell transcriptome from heterogenous populations, we went on to directly study microbial single-cells in tissue samples, which has been impossible with other methods. After gavage of mice with Cy3-amino-D-alanine (Cy3ADA) and Fam-amino-D-alanine (FADA), various morphologies of fluorescent bacteria were observed in intestinal lavage fluid, indicating that intestinal bacteria can be specifically labeled with amino-D-alanine (Fig.5a). Therefore, we further observed fluorescent bacteria near the mucus of a mouse colon and inside host cells (Fig.5b). Single bacteria could be extracted from these regions either in a fluorescence-dependent or -independent manner (Fig.5c and d). We selected 25 bacteria from three fields for genome sequencing, obtaining 3 high-quality and 5 medium-quality SAGs. Among these bacteria, there are typical mouse gut bacteria like *Prevotella copri A*, *Escherichia flexneri*, *Turicimonas muris*, *Bacteroides intestinalis*, *Bacteroides dorei* as well as more aerobic and peritoneal species such as *Cutibacterium acnes*, *Staphylococcus capitis* (Fig.5e)^40, 41^. These results suggested that LIFT could be applied to tissue samples.

**Figure. 5.**
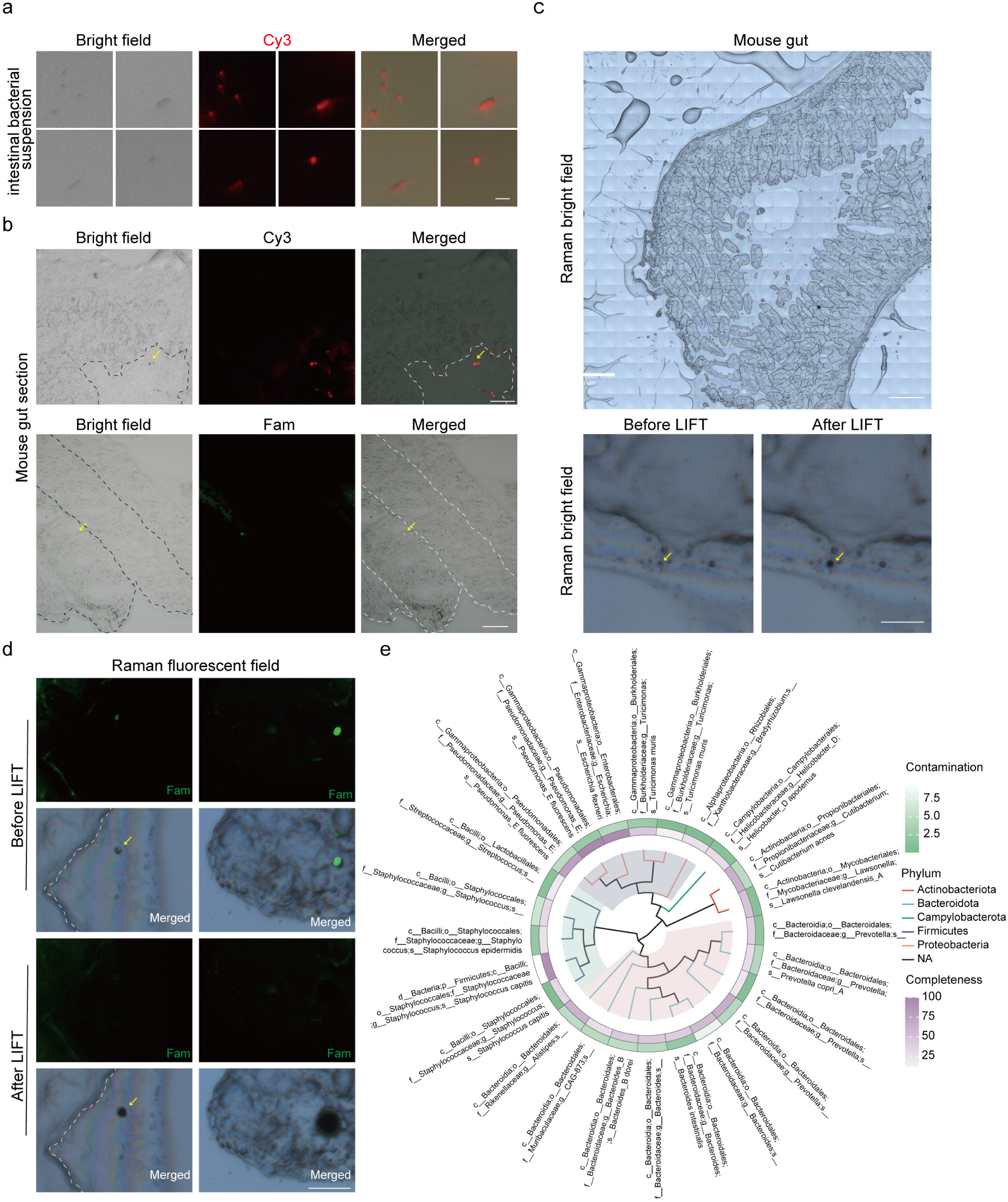
LIFT ejection of bacteria in mouse gut in bright and fluorescent field. **(a)** Representative images of bacterial morphology in mouse large intestinal lavage suspension after gavage with Cy3-ADA in mice. Scale bar, 10μm. **(b)** Cy3-ADA and Fam-ADA labeled bacteria on mouse large intestine sections. Scale bar, 100μm. **(c)** Localized composite image of mouse large intestinal tissue sections (upper panel). Scale bar, 200μm; LIFT ejection of bacteria on mouse large intestine sections in bright field (lower panel). Scale bar, 10μm. **(d)** LIFT ejection of bacteria on mouse large intestine sections in fluorescent field. Scale bar, 10μm. **(e)** 25 single bacteria were isolated from three fields of view in the mouse large intestinal tissue sections. A phylogeny constructed from 25 obtained SAGs genome is represented by the inner dendrogram. The phylum of each SAGs is indicated by the branch color. Completeness and contamination of each SAGs are represented by two heatmaps overlay on the phylogenetic tree. The species name of each SAGs is listed in the outermost ring.

We next applied the method to tumor samples from human patients. Resected tumor tissues from a patient diagnosed as colorectal cancer (Sup. Fig. 9a) were cut into tissue sections and loaded onto the slide for microbial single-cell omics (Fig.6a). We confirmed cancer infiltrating and staging by immunohistochemical staining of marker genes such as P53, BrafV600E, Ki67 (Sup.Fig.9b). Using antibodies against bacterial lipopolysaccharide (LPS) and vancomycin to detect Gram-negative and Gram-positive bacteria respectively, we observed microbes with diverse morphological features including rod shape and spherical shape, distributed in colorectal cancer (Sup. Fig. 9c). Without special treatment, we searched for microbes in the bright field. The microbes have distinct Raman spectral fingerprints compared to histiocytic Raman spectrum (Sup. Fig. 9d), which means this technology can very effectively avoid sequencing of host cells or collection of non-microbial particles. For this colorectal cancer sample, a 1.33μm long and rod-shaped single cell was spotted and separated for genomic sequencing combined with Raman spectrum acquisition (Fig.6b and c). We obtained a medium-quality SAG identified as *Bacteroides intestinalis* based on Mash analysis (Fig.6d). Phylogenetic analysis of all *B. intestinalis* genomes from metadata-gut of Mash database and our SAG revealed that SAG was similar to MGYG000028343 which is also from China (Fig.6e).

**Figure. 6.**
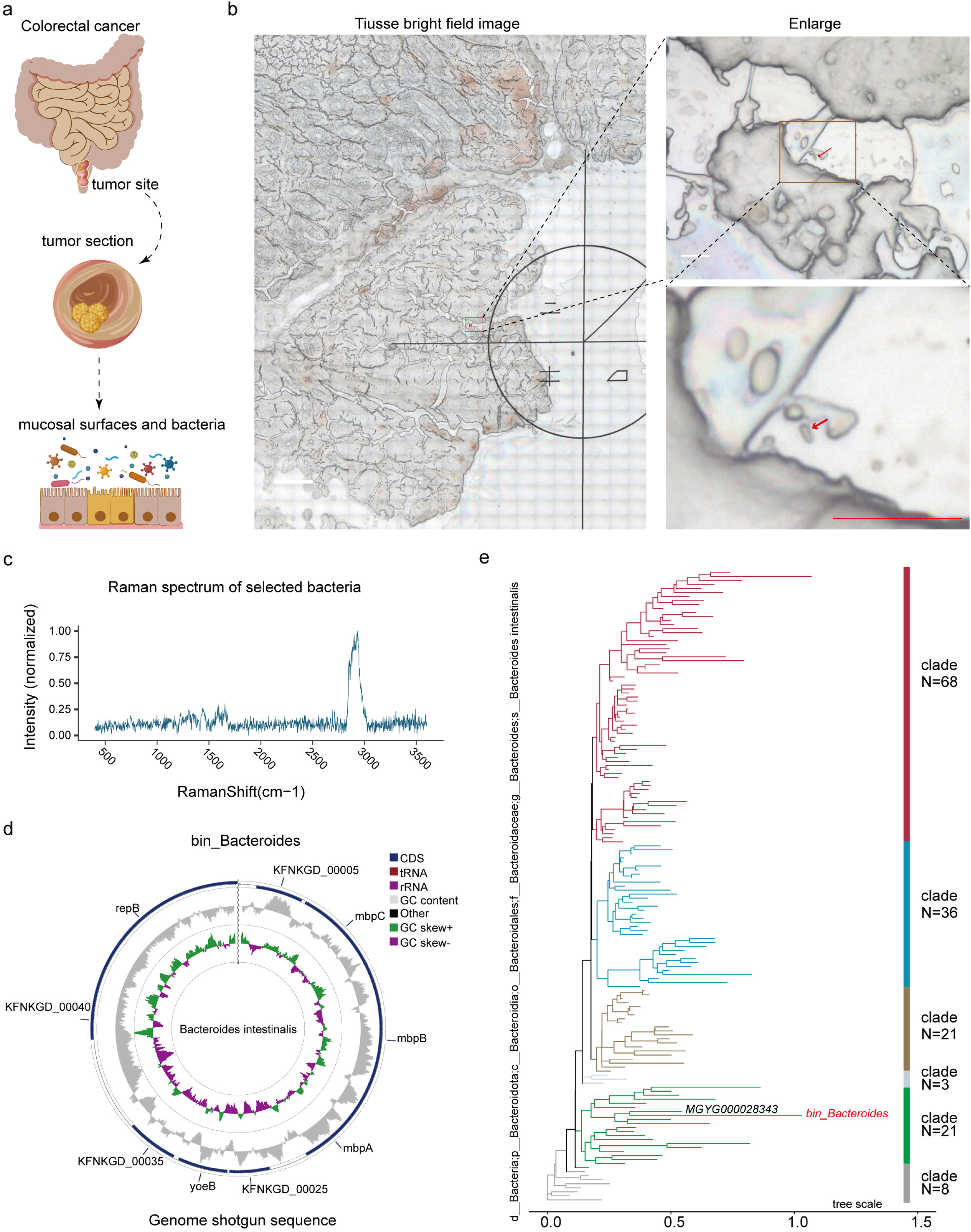
Genomic sequencing for single microbe in tissue. **(a)** Diagram for the colorectal cancer sample. **(b)** Tissue section containing the microbial single-cell ejected and genome sequenced. Scale bar, 10μm. **(c)** Raman spectrum of the indicated single cell. **(d)** Visualization of obtained SAG (bin Bacteroides) by CGView tool displaying GC skew, feature labels, GC content, divider rings, and tick density. **(e)** Phylogenetic tree was constructed by obtained SAG (bin Bacteroides) and all *Bacteroides intestinalis* genomes downloaded from metadata-gut of Mash database. Among Bacteroides intestinalis genomes from Mash database, genome *MGYG000028343* was isolated from China.

Moreover, a rod-shaped single cell adhered to intestinal mucosal was separated for RNA sequencing (Fig.7a). After removal of host gene reads, we obtained de novo transcriptome assembly identified as *Bacteroides xylanisolvens* by GTDB-Tk, in which 70% contigs and 75% transcripts (2756 transcripts) belonged to *Bacteroides* (Fig.7b). Of note, we observed dust particles in the lungs of this colorectal cancer patient with 50 years of smoking when cancer metastasized to the lungs (Sup. Fig. 10a and Sup. Fig. 10b), which is consistent with the finding that *Bacteroides xylanisolvens* is involved in nicotine degradation in smokers ^42^. Among the ∼3616 genes expressed were nicotine-degrading gene nicX, membrane proteins SusC/RagA family TonB-linked outer membrane protein, ABC transporter, sialic acid-specific acetylesterase and porin (Fig.7c), which are crucial for nicotine degradation, nutrients uptake and thrive of *Bacteroidetes* in the gut ^43-45^. Gene ontology analysis further verified that these genes were enriched in pathways of passive transmembrane transporter activity, sialic acid transmembrane transporter activity and porin activity (Fig.7d). These results suggest the application potential of this method for tissue microbe detection *in situ* and single cell spatial microbiome.

**Figure. 7.**
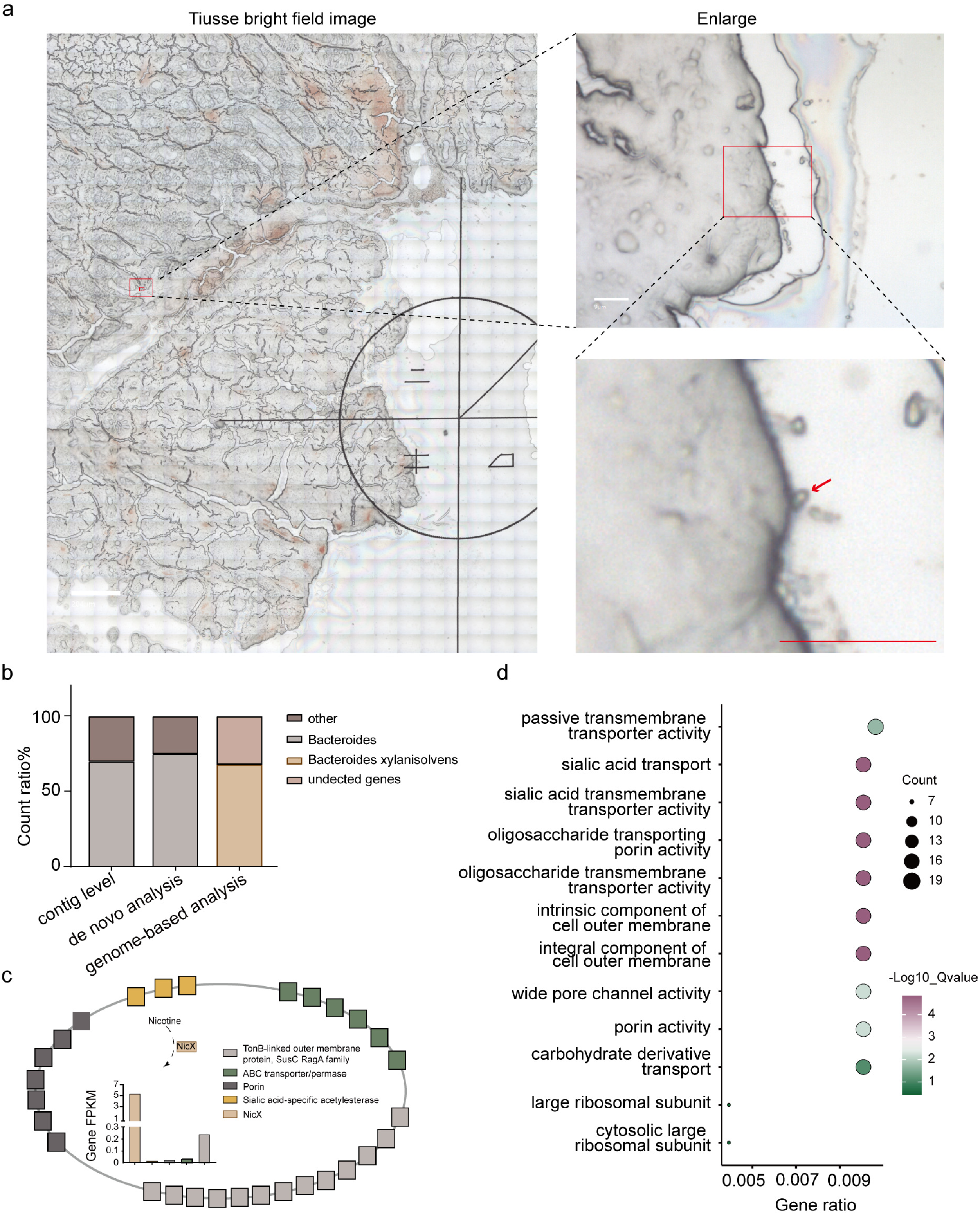
Transcriptome sequencing for single microbes from tissue. **(a)** Tissue section as in Figure 6, with inlets indicating the position of the bacteria for scRNA analyses. Scale bar, 10μm. **(b)** Count ratio of contigs, transcripts in single cell assembled transcriptome and genes by genome-based analysis. Contig classification was annotated by Kraken2. Transcript annotation was referred against de novo transcriptome annotated by eggnog-mapper software. Gene annotation was referred against *Bacteroides xylanisolvens* genome (total 5306 genes). **(c)** The detection of nicotine degradation gene nicX and proteins located on cell membrane. Gene FPKM was calculated by Bakta. **(d)** Gene ontology analysis of transcripts classified to *Bacteroidetes*.

## Discussion

Our study provides a method based on LIFT technique for cell isolation and further high-quality single cell sequencing. This method can precisely and automatically isolate individual bacteria from complex samples, either in a label-dependent or label-free manner. After rapid automated sorting, the cells were transferred to an automated platform for nucleic acid extraction and amplification, in preparation for subsequent single-cell genome or transcriptome sequencing to enhance throughput. We are positive that further reducing the reaction volumes, optimizing the reagents, the plasticware and the precision selection process will further demonstrate the advantage of this single-cell omics approach that does not rely on microfluidics and exhaustive sequencing of abundant cells.

Knowing the complete single-cell genome and transcriptome would be very useful to understanding fundamental questions of the microbiome beyond model organisms. Both the single-cell genome and the transcriptome could be either reference-based or de novo assembled, the latter means applicability to any new sample instead of a model bacterium. As is commonly seen for single-cell studies of mammalian cells, merely increasing the number of cells sequenced by a few fold would not necessary ensure identification of rare populations. With our approach, rare populations could be enriched before sequencing. Moreover, our method provided multidimensional information for the study of single cell microbe, including bacterial shape, size, and Raman spectrum. Visually, the existing software could already be set to select for cell size, shape, area, etc (Sup. Fig.5b). Further standardization and algorithm development for the single-cell Raman spectra would allow more precision in sub-species analyses.

Another major advantage of this LIFT-based single-cell approach is that complex samples require minimum pretreatment, and can contain other particles or other cells. The separation of neighboring cells would also allow elucidation of microscopic cooperation in a native environment in future studies. We have previously reported multiple *Bacteroides* species in an Austrian cohort of colorectal adenoma and carcinoma ^46, 47^ and *B. fragilis* were typically studied in mice models of colitis ^48^. Here the scRNA analyses have tentatively identified metabolism of nicotine, mucin, etc. We have not yet tried FFPE samples, which should be theoretically possible, so long as the nucleotides have not been too degraded. Further application of our microbial single-cell omics method to clinical samples would facilitate patient stratification and treatment.

In summary, here we present an optics-based strategy for precision single-cell microbial genomics, and transcriptomes that could fully capture the genomic landscape of host-associated microbiome, and allow nucleotide-resolution understanding of microbial gene transcription and turnover in individual cells in their native environment.

## Methods

### Animals

Specific pathogen-free C57BL/6 mice (female, 8 weeks old) were purchased from BESTEST Biotechnology Co., Ltd (Zhuhai, China) and maintained in the laboratory Animal Center of Greater Bay Area Institute of Precision Medicine (Guangzhou, China). These mice were bred in a temperature-controlled environment (25℃, 12-hour light/dark cycle) with free access to food and clean drinking water. All animal experiments were approved by the Animal Ethics Committee at Greater Bay Area Institute of Precision Medicine and were performed in accordance with guidelines approved by the Institutional Review Board of Fudan University.

### Study participants and sample collection

All the adult volunteers were not known to have chronic diseases, and did not take antibiotics within the past three months. All healthy volunteers and the patient had written informed consent. All the samples were self-collected into a sterile tube, and handed in for further processing on the same day. The study was approved by the Institutional Review Board of Fudan University. Tissue collection was approved by the Ethics Committee at The Affiliated Panyu Central Hospital of Guangzhou Medical University and was conducted following guidelines established by this committee.

### Laser Induced Forward Transfer (LIFT) of bacterial cells

The equipment, PRECI SCS-R300 (Hooke Instruments, Ltd., China), was turned on and adjusted according to standards following the manufacturer’s handbook *^49^* (Sup. Movie 1). The optical setup is shown in Sup. Fig. 1. Laser used for LIFT was nanosecond single pulse laser with 532nm wavelength, 5ns pulse width and 3.4μJ energy, which could produce a single adjustable spot capable of ejecting 0.5-20μm microorganisms. For complex shape and larger microbes (greater than 20μm), LIFT combined with beam shaping could successfully sort them. Laser for Raman spectral acquisition was continuous laser with a maximum laser power of 50mW, 532nm wavelength. The parameters of the laser for the acquisition of Raman spectra are set to 3mW, 3s.

The filtered and washed samples were spotted onto a quartz slide coated with 25nm thickness of aluminum film (HSC24, Hooke Instruments, Ltd.), and air-dried in the hood. The Hooke chip was designed to incorporate a thermal insulation structure to effectively protect the cell from heat-induced damage. Visually selected bacterial cells (the manufacturer’s software) were LIFT-transferred into collectors (HSR04, Hooke Instruments, Ltd.) using a laser energy below 75nJ (exceeding the minimum energy required by most microbes), and then flipped into individual PCR tubes with brief spinning at 2000g (Yooning Mini-6K).

### Whole genome amplification and sequencing from single bacterial cells

Following cell lysis at 65℃ for 10 min, Multiple displacement amplification (MDA) was performed using the REPLI-g scDNA Polymerase, according to kit instructions (REPLI-g single cell kit, #150345, QIAGEN). Sequencing libraries (PE150) were constructed without PCR, and sequenced on a DNBSEQ-T7 high-throughput sequencer (MGI).

### Genome assembly and analyses

Quality control on paired-end sequencing reads of raw data was performed by fastp (0.23.2) tool. The filtered reads were assembled by metaSPAdes (3.13.0) in single cell mode. Based on binning information provided by Kraken2 (2.0.7), the obtained assembled bins were processed with bin refinement using BinSpreader. The quality of genome bins was analysed by Checkm2 and QUAST. GTDB-Tk (1.0.2) was used to classify and taxonomically assign genome bins. Phylogenetic trees were constructed by PhyloPhlAn (3.0.67) tool.

### Whole transcriptome amplification and sequencing from single bacterial cells

In addition to phosphate buffered saline (PBS) and lysis buffer according to the QIAGEN kit (#150065), cell lysis was performed at 24℃ for 5 min, in the presence of murine RNase Inhibitor (Vazyme), and MetaPolyzyme (MAC4L, Invitrogen). Total RNA Reverse-transcription, ligation and MDA were performed using the REPLI-g WTA single cell kit (#150065, QIAGEN), with extended reaction time for bacteria instead of mammalian cells: 2 hours for ligation, 4 hours for MDA. Sequencing libraries (PE150) were constructed without PCR, and sequenced on a DNBSEQ-T7 high-throughput sequencer (MGI).

### Single-cell transcriptome analyses

For de novo transcriptome analysis, filtered reads of scRNA sequencing sample was assembled using rnaSPAdes (3.15.5)^50^. Bakta (1.8.1) conducts a comprehensive annotation of assembled transcriptome. Protein sequences generated by Bakta (version 1.8.1; AMRFinderPlus for AMR gene annotations; a generalized protein sequence expert system for identity, distinct coverage and identity and priority values for each sequence) were further annotated by eggnog-mapper (5.0.2) for gene function. KofamScan is used to annotate protein sequences with KEGG Orthology (KO). DeepFRI is used to predict Enzyme Commission (EC) numbers and Gene Ontology (GO) terms for Bakta annotated genes. For analyzing the distribution and characteristics of oligoA, bioawk tool is used to analyze oligoA positions in gene/contig level, and then process the results to obtain matrix containing gene/contig lengths, oligoA positions, gene start and gene end positions. All transcripts in de novo transcriptome were evenly divided into ten percentile according to gene length and the number of occurrences of oligoA positions within each percentile range of the gene lengths was recorded. Standard deviation of oligoA distribution was calculated according to the fraction of counts in each percentile range of the gene length. The proportion of oligoA occurrence was normalized by R package “Mfuzz”. The analysis of poly(A) with nine adenylate residues (9A) or more followed these steps. Bowtie2 is used to align reads to transcripts and calculate gene FPKM values. For genome-based analysis, reads are referred against *Bacillus licheniformis* genome and genome annotation GTF file (GCF_002074095.1) downloaded from NCBI database. After alignment with Bowtie2, gene quantification was performed using featureCount (subread2.0.3). Genes expression with fold change ≥1.5 and pvalue < 0.05 were determined as significant change using DESeq2 (version 1.20.0). Gene enrichment of Biological Process (BP), Molecular Function (MF), Cell Component (CC) was analysed by R package Clusterprofiler with pvalueCutoff = 0.05. Heatmaps of gene expression were using R package “pheatmap”. The bar plots, bubble plots of GO analysis, scatterplots, and boxplots were visualized using on-line tool imageGP (http://www.ehbio.com/ImageGP/index.php/Home/Index/index.html).

### *Bacillus licheniformis* culture and transmission electron microscopy

The stored *Bacillus licheniformis* strain was inoculated into the Columbia Broth medium overnight at 37°C. 100μl *Bacillus licheniformis* suspension was transferred to 2ml medium containing 10% Columbia Broth, in which PH was adjusted to 4.0 by adding citric acid. *Bacillus licheniformis* were cultured in citric acid-amended 10% Columbia Broth for day 3 (exhibited a variety of morphologies) and day 9 (almost all transformed into spore at day 9). Then *Bacillus licheniformis* suspension was passed through 40μm cell strainers to eliminate cell clumps. The resulting filtrate was centrifuged at 12,000g for 3 minutes to collect bacterial pellets. These pellets were then washed three times with 1.5 ml of PBS each and finally resuspended in 0.7ml ddH2O. Prior to using PRECI SCS R300 devices, 1-2 µl of the cell suspension was dropped onto chip and allowed to dry either by evaporation at room temperature or by exposure to sterile airflow in a biosafety cabinet.

For transmission electron microscopy, citric acid treated *Bacillus licheniformis* were fixed with 4% PFA and stored at 4℃. The copper mesh was incubated in 10μl fixed *Bacillus licheniformis* solution for 2 minutes. After removal of excessive solution using filter paper, copper mesh was incubated with 1% phosphotungstic acid (PTA) for 2 minutes and observed by electron microscope (FEI Tecnai G2 spirit).

### Metabolic labeling with fluorescent D-alanine in culture

TADA (tetramethylrhodamine-amino-D-alanine), Cy3ADA (Cyanine3-amino-D-alanine) and FamADA (5-Carboxyfluorescein-amino-D-alanine) were purchased from GuoPing Pharmaceutical (Anhui province, China). *Staphylococcus capitis*, *Streptococcus oralis*, *Lactobacillus plantarum*, *Akkermansia muciniphila* and *Clostridium butyricum* were isolated from faeces, urine and saliva using our LIFT technology and cultured in Gifu Anaerobic Medium at anaerobic incubator. The *Sphingomonas sp000797515* were generously provided by HOOKE Instruments Ltd. The *Sphingomonas sp000797515* were cultured in Luria-Bertani Medium and incubated with TADA probes (40mM) at 37℃ for 3 hours. After incubation, the *Sphingomonas sp000797515* were washed with PBS thrice, and mixed with other single-strain bacteria in ddH2O for the following analyses. The *Sphingomonas sp000797515* genome provided was assembled from our single strain suspension sequencing.

### Metabolic labeling with fluorescent D-alanine in mice

Cy3ADA and FamADA were administered to mice via gavage to label mouse gut microbiota. In brief, Mice were gavaged with Cy3ADA and FamADA probes dissolved in 200μl of PBS at 1 mM concentration. 6 hours later, Mice were anesthetized using sodium pentobarbital (50 mg/kg). A part of mouse large intestine was excised and minced using scissors in 2ml of PBS buffer. Subsequently, the minced tissue was passed through 40μm cell strainers to eliminate nonbacterial tissues and food debris. The resulting filtrate was centrifuged at 12,000g for 3 minutes to collect bacterial pellets. These pellets were then washed three times with 1.5ml of PBS each and finally resuspended in ddH2O. The bacterial suspension was placed under a Leica fluorescence microscope (ECLIPSE Ni-U) to observe the labeling of the bacteria.

### Tissue pretreatment for LIFT cell sorting and Hematoxylin-Eosin staining, Immunofluorescence staining, Gomoris staining

After euthanizing the mice, a small segment of colon tissue (0.5-0.8cm length) free of food debris was excised, embedded in OCT (optimal cutting temperature compound), and frozen. The frozen colon tissue was sectioned into slices with 6μm thickness and mounted flat onto glass slides and LIFT chip.

Several pieces of 2mm thickness tissues (20×20×2mm) separated by laparoscopic colorectal cancer surgery were placed on filter paper and dry gauze pad. After removal of excessive water, tissues were transferred to supporter of freezing microtome (Leica CM1860 UV) and embedded with OCT. In order to ensure the operation of the tissue section, OCT embedded tissue was placed on cryo-stage until it’s a solid ice cube. Processed tissue was sectioned into 7μm thickness for Raman spectrometry and LIFT on PRECI SCS-R300 (Hooke Instruments, Ltd., China) with 75nJ laser energy.

Dehydrated and paraffin embedded tissues were cut into 5μm thickness and were stained with hematoxylin-eosin after de-paraffinization. For immunofluorescence staining, tissue sections were treated with antigen retrieval buffers, and then permeabilized and blocked by mixed solution of 0.3% Triton X-100 and 5% BSA for 1 hour. Primary antibodies were used to incubate tissue sections overnight at 4°C. Phosphate Buffer Saline containing 0.1% Tween-20 (PBST) was used to wash tissue sections five times. Well-washed sections were carefully incubated with secondary antibodies for 1 hour at room temperature. DAPI (4’,6-diamidino-2-phenylindole) diluted at PBS with concentration 1:1000 was used to cell nuclei staining. Immunofluorescence images were captured by Leica sp8 system. Antibodies and their working concentration are listed as followed: anti-CK20 (ZSGB-BIO, #ZA-0574, 1:200), CDX2 (ZSGB-BIO, #ZA-0520, 1:200), Villin (ZSGB-BIO, #ZA-0575, 1:300), MSH2 (ZSGB-BIO, #ZA-0622, 1:250), MSH6 (ZSGB-BIO, #ZM-0367, 1:250), MLH1 (ZSGB-BIO, #ZM-0152, 1:300), PMS2 (ZSGB-BIO, #ZA-0542, 1:200), P53 (ZSGB-BIO, #ZM-0408, 1:200), BrafV600E (Roche, #760-5095, 1:300), and Ki67 (ZSGB-BIO, #ZM-0166, 1:300), anti-LPS (Cloud-Clone, #MAB526Ge22, 1:150), anti-Vancomycin (Invitrogen, #V34850, 1:150).

Gomoris staining method followed the manufacturer’s protocol (Baso diagnostics. lnc. Zhuhai. China, #BA4375A). Briefly, 5μm thickness paraffin tissue sections were brought to water via xylene and descending grades of ethanol for deparaffinization. Following nuclear staining by Weigert’s Iron Hematoxylin, sections were placed in Solution A for 15 minutes. After rinsing off the stain with distilled water, sections were stained with Solution B. After dehydration with ethanol and clearing with xylene, sections were mounted with a resinous medium.

## Supporting information

supplemental vedios

## Acknowledgements

The authors were grateful to Zhong Yang, Lili Li, Jiong Ma, Xu Zhang, Hang Li, Meiya Lin for equipment support. Ping Sun, Miaoyi Kuang for sequencing support. We thanked Prof. Hao Chen (Shanghai Institute of Materia Medica, Chinese Academy of Sciences) and Prof. Xin Feng (Jilin University) for their suggestions on mouse gut microbiota labeling.

## Author contributions

Xihong Lan and Huijue Jia conceived and designed the methodology; Xihong Lan developed and performed the experiments with assistance from Jiayi Wu; Xihong Lan and Jiayi Wu optimized sequencing library preparation; Jinhua He and Xiaoying Zhang prepared tissue sections; Xihong Lan, Qiaoxing Liang and Ruidong Guo performed the sequencing data analysis; Xihong Lan wrote the original manuscript; Huijue Jia and Guoping Zhao revised the manuscript; Huijue Jia supervised the study.

## Correspondence and requests for materials

Further information and requests for resources and reagents should be directed to and will be fulfilled by the lead contact: Huijue Jia

