## supplemental figures for "Microbial single-cell omics in situ"

### Supplementary Information

**Supplementary Fig.1 to Fig.10**

**Movies S1 to S4**

**Tables S1 to S3**

Supplementary Fig.1

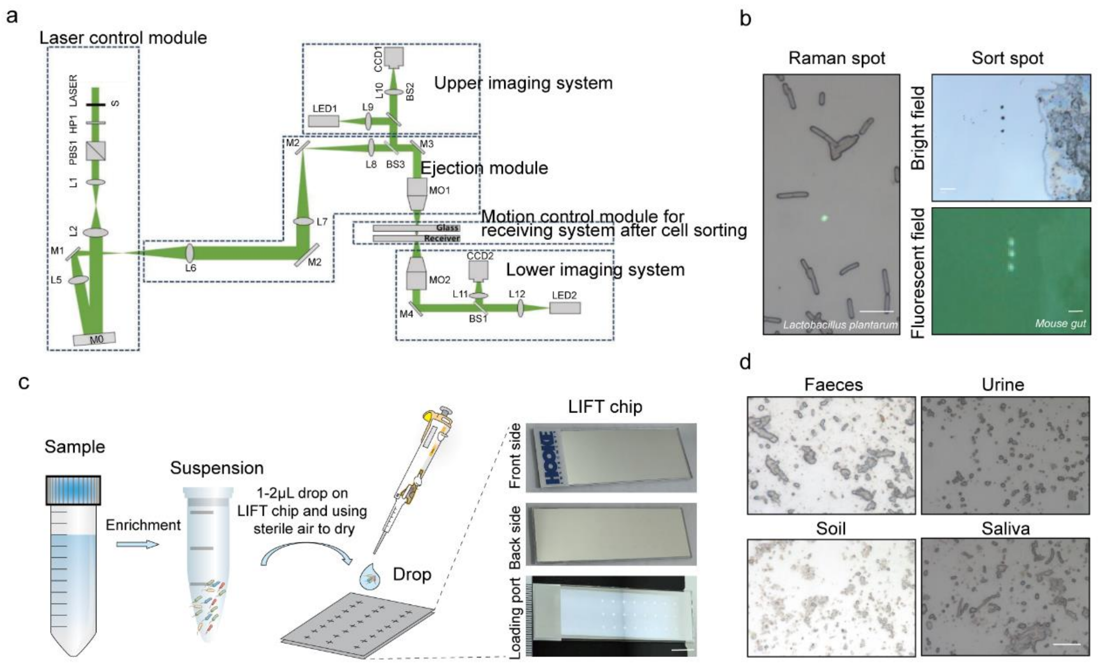


**Supplementary Fig.1 Optical diagram for LIFT, imaging and Raman spectral acquisition, and sample pre-processing steps before LIFT. (a)** Our PRECI SCS R300 devices consist primarily of laser control module, imaging system, ejection module and motion control module. The laser control module was used for regulation of laser energy and power, the beam expanding and on-off control. The upper imaging system was mainly used for imaging the laser ejection spot to calibrate the position. The lower imaging system was mainly responsible for the imaging of microbial cells on the chip and the receiver. The ejection module was used to focus the laser spot and eject cells. The motion control module control the chip motion and coordinated with the imaging system to precisely receive the ejected cells. S: Shutter, HP: Half wave plate, L: Lens, M: Mirror, BS: Beam splitter, PBS: Polarizing beam splitter, MO: Microscope objective. **(b)** Spot of laser for Raman spectral acquisition (left panel); LIFT ejection (70nJ) resulted in shedding of thin metal coating along with the cell, which formed black spot (bright field) and bright holes observed under fluorescent field in mouse gut chip (right panel). **(c)** Workflow for the minimum sample treatment before imaging and LIFT ejection, the slide contains 24 plus sign markings for sample loading (left panel); The back side of HOOKE LIFT chip contains 24 sample loading ports (right panel). **(d)** Without the need for complex steps, LIFT is suitable for microbial single cell isolation from multiple samples including faeces, urine, soil and saliva, etc. Various forms of microbes from these samples can be clearly observed after treatment.

Supplementary Fig.2

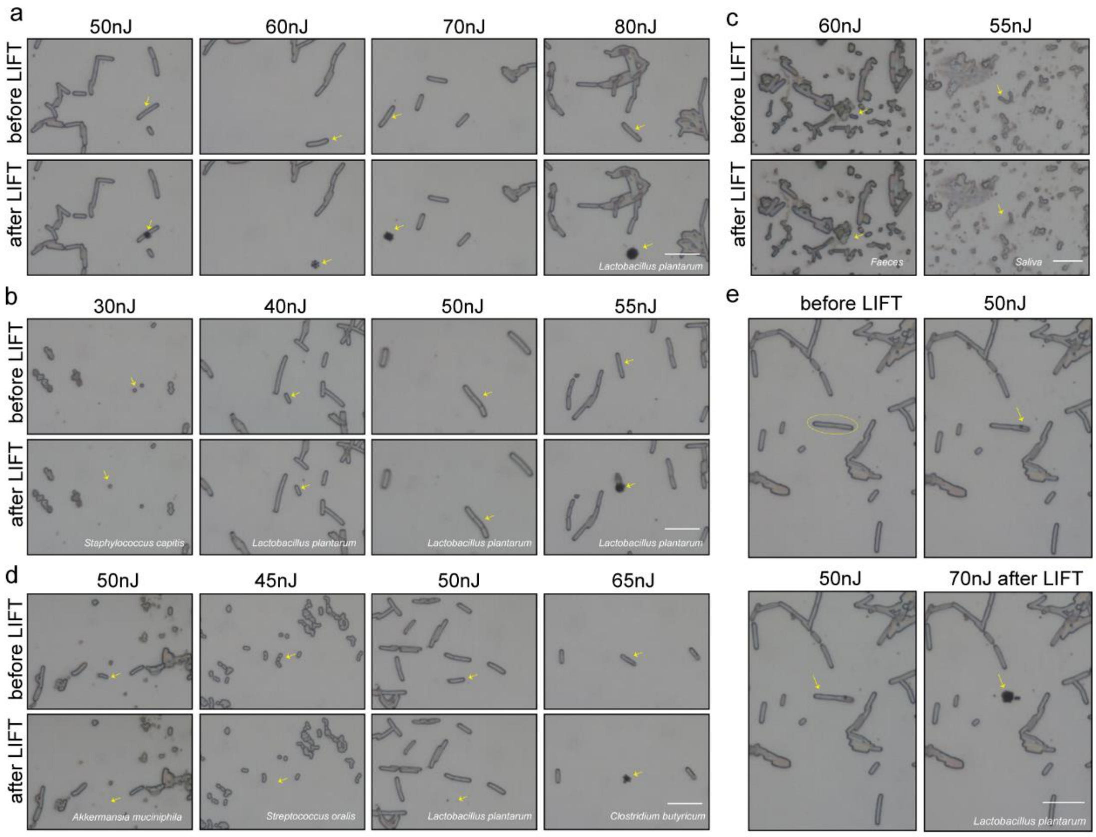


**Supplementary Fig. 2 Laser energy is essential for the successful sorting of microbial single cells. (a)** The same size of *Lactobacillus plantarum* were ejected at different laser energy showing that cells could be successfully ejected when the energy exceeds the minimum energy required for cell sorting and the spot of the laser used for LIFT ejection become larger as the energy increased. Scale bar, 10μm. **(b)** Less energy than required resulted in cell shifting (*Staphylococcus capitis* 30nJ, *Lactobacillus plantarum* 40nJ), cell ejection failure (failed ejection of *Lactobacillus plantarum* 50nJ) and cell rupture (fragment of *Lactobacillus plantarum* 55nJ). Scale bar, 10μm. **(c)** When energy is suitable, LIFT can precisely isolate microbial single cells from complex samples such as faeces and saliva even with densely packed bacteria. Scale bar, 10μm. **(d)** Different microbial cells such as *Akkermansia muciniphila*, *Streptococcus oralis* occurring in short chains, *Lactobacillus plantarum* and *Clostridium butyricum* required different levels of LIFT energy. Scale bar, 10μm. **(e)** In addition to increasing cell LIFT energy, larger size *Lactobacillus plantarum* could be ejected by multiple LIFTs as indicated 50nJ at left, 50nJ at right and then 70nJ in the middle of the cell. Scale bar, 10μm.

Supplementary Fig.3

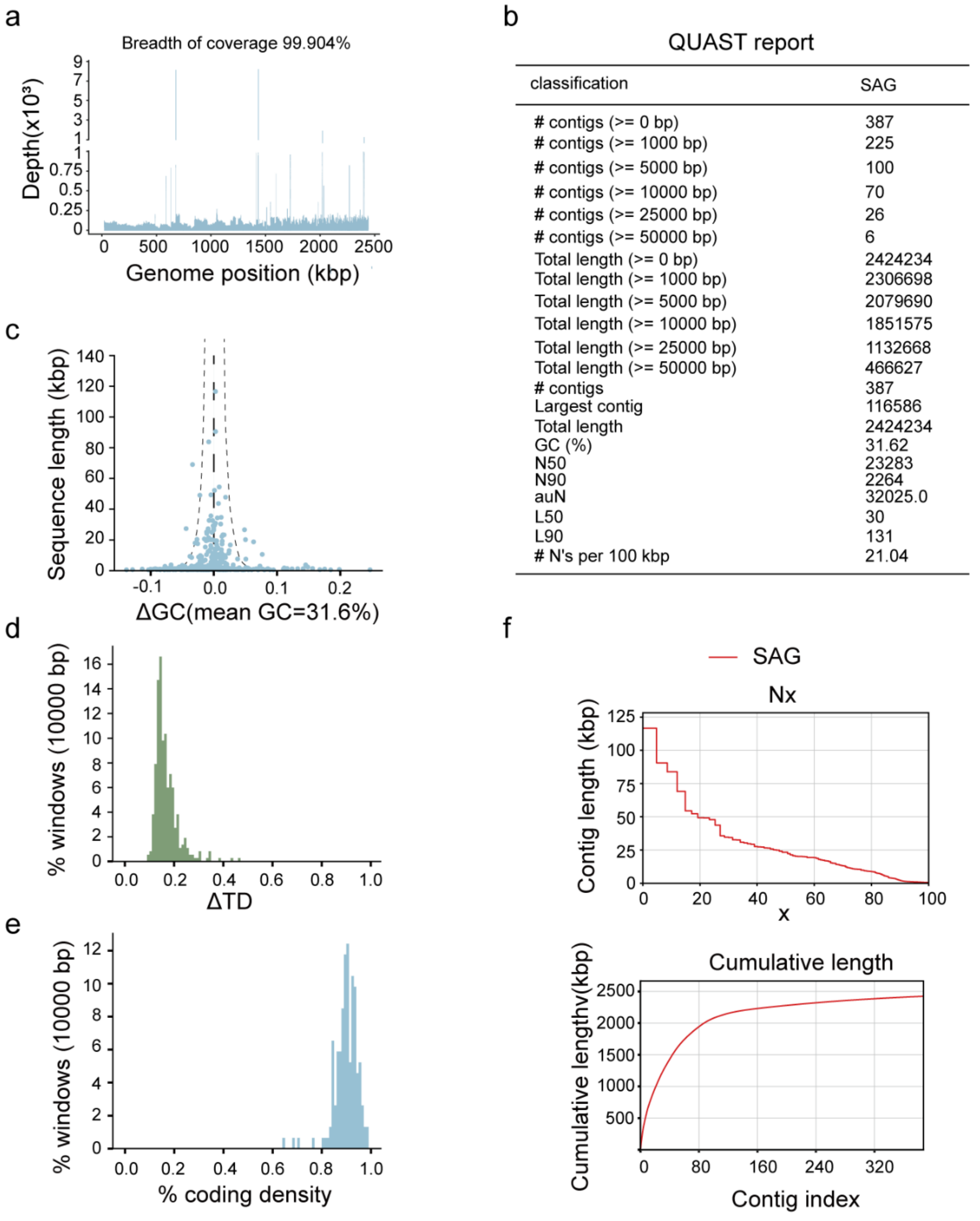


**Supplementary Fig.3 Evaluation of single-cell assembled genome quality. (a)** Calculating the per-base depth and breadth of coverage in the single-cell assembled genome. **(b)** The SAG (the obtained single-cell assembled genome) genome quality by QUAST analysis. **(c)** GC content distribution of sequences within bin SAG by CheckM. **(d)** Tetranucleotide signature distribution of SAG by CheckM indicate low contamination. **(e)** Coding density distribution of sequences within SAG. **(f)** Contig length distribution of SAG visualized by Nx plot (upper panel); the cumulative length plot indicates the presence of longer contigs in SAG (lower panel).

Supplementary Fig.4

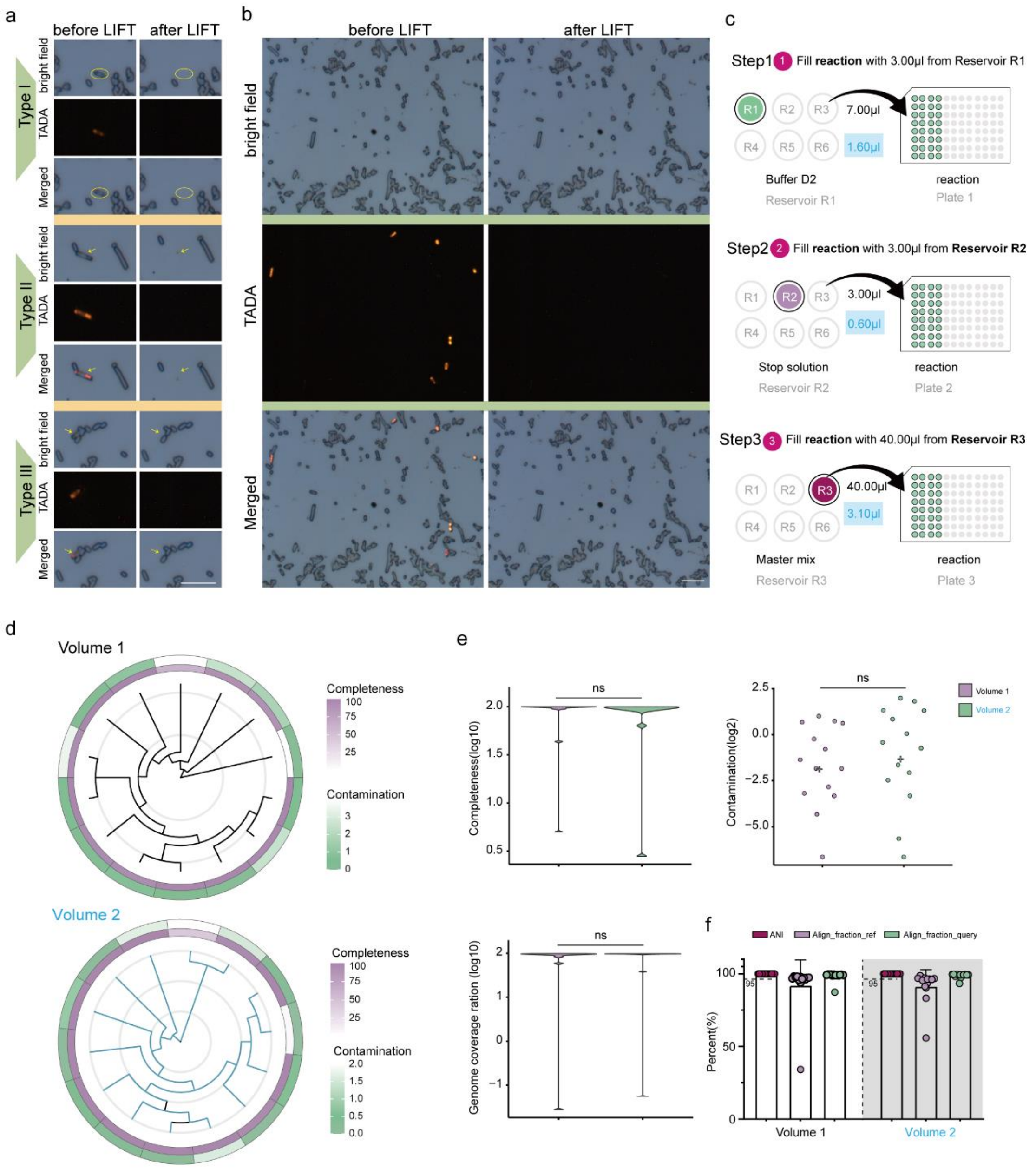


**Supplementary Fig.4 Automated isolation of single TADA-** ***Sphingomonas sp000797515*** **cell from a mixture of multiple bacteria and attempt to reduce reaction volume. (a)** Precise LIFT ejection of single TADA-*Sphingomonas sp000797515* mixed with *Bacteroides fragilis*, *Lactobacillus plantarum* and *Clostridium butyricum*. The bacteria were extracted from sparse region (type I) and densely packed region when two bacteria were very close (type II) and partially overlapped (type III). Scale bar, 10μm. **(b)** Automated LIFT ejection of single TADA-*Sphingomonas sp000797515* in fluorescent modality. Representative images before and after LIFT ejection. Scale bar, 10μm. **(c)** A designed pipetting program for Firefly including adding lysis buffer, adding stop solution and adding master mix automatically. After cell sorting, an attempt was made to reduce the reaction volume to one-tenth of the original volume. **(d)** 28 cells from a single field of view were evenly divided and used to react with two different volumes for sequencing, yielding 28 single amplified genomes (SAGs). The two phylogenetic trees constructed from the 14 *Sphingomonas sp000797515* SAGs (Twelve high-quality SAGs, one medium-quality SAG, one low-quality SAG) were respectively represented by two inner dendrograms. Completeness and contamination of each SAGs are represented by two heatmaps overlay on the phylogenetic tree. **(e)** The quality of the SAGs obtained from the two different reaction volumes was compared, including completeness, contamination and coverage. **(f)** The ANI of SAGs obtained by two reaction volume against reference genomes were analyzed using the skANI software. The ANI values of medium- and high- quality SAGs are both over 95%. Align fraction ref: Reference sequence alignment coverage. Align fraction query: Query sequence alignment coverage. Data are represented as mean ± SE (ns: P >0.05, *P < 0.05, **P < 0.01, ***P < 0.001).

Supplementary Fig.5

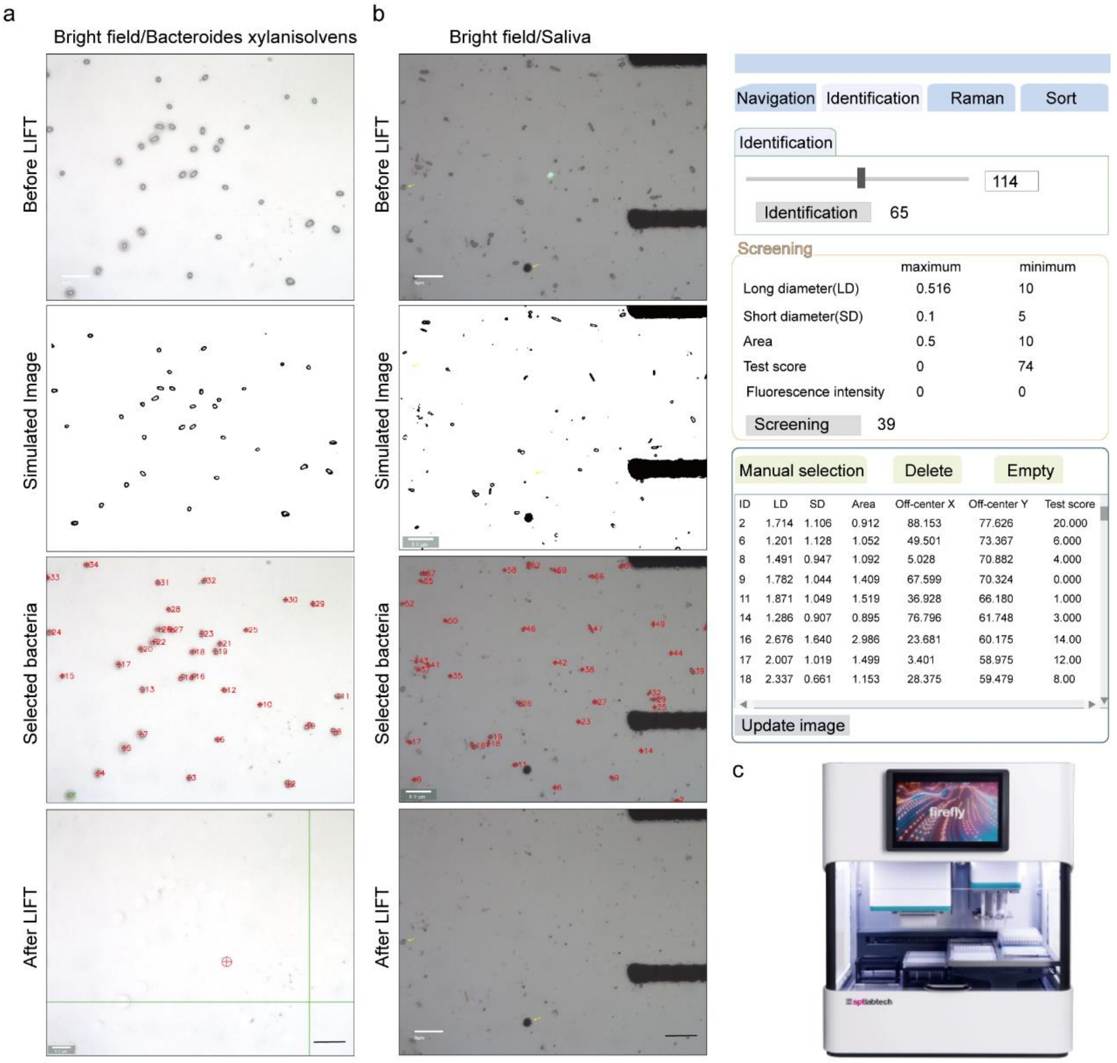

**Supplementary Fig.5 Automated single-cell ejection in single bacterial suspension and complex sample (salvia). (a)** Automated single-cell ejection for single bacterial suspensions. All automatically tagged *Bacteroides xylanisolvens* were completely ejected. **(b)** Automated single-cell ejection for bacteria from complex sample salvia. When microbes were loaded onto slide, simulated image could be obtained by adjusting the value of Identification (right panel) to cover most cells, and then these cells were numbered. Adjusting the value of the parameter Long diameter (LD), Short diameter (SD), Area, Test score and Fluorescence intensity could help to select single cells more accurately. Debris and black spot (indicated by yellow arrow) can be excluded manually. Those selected or unselected cells that are particularly discrete could be delete or added by pressing button Delete and Manual selection. The information on the selected cells was listed by cell ID. **(c)** Automated molecular experiment platform Firefly (SPT Labtech).

Supplementary Fig.6

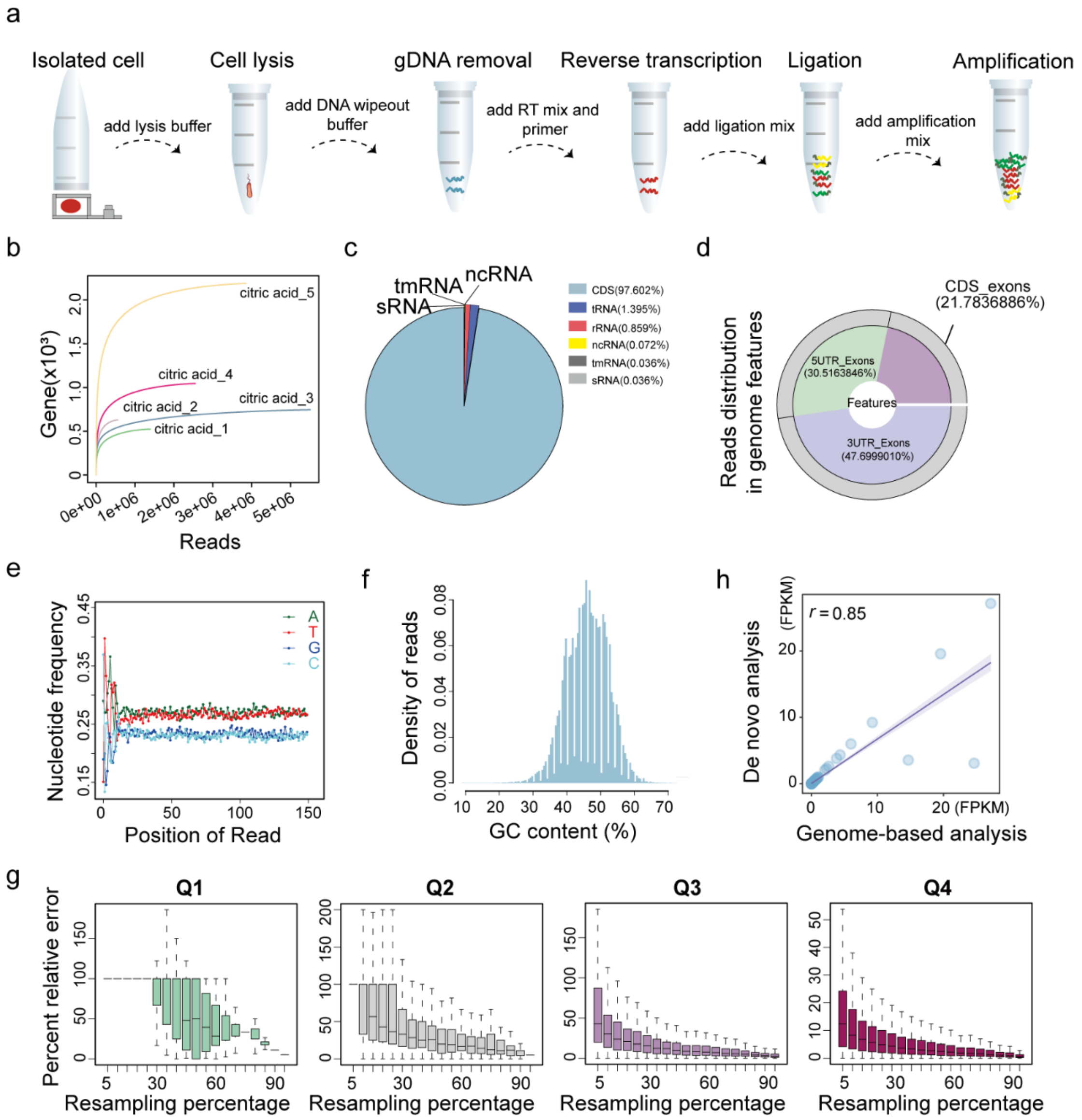


**Supplementary Fig.6 Quality control and visualization of representative scRNA-seq data. (a)** Diagram illustrating steps for single cell RNA reverse transcription and amplification. **(b)** Rarefaction curve of samples treated with (citric acid_1/2/3/4/5) citric acid indicated sufficient sequencing depth. The mapped genes selected for rarefaction curve were protein coding genes against reference genome, which varied from less to more in each sample. **(c)** Gene type annotation of a representative sample in pie chart. **(d)** Reads distribution over reference genome feature including CDS exon, 5'UTR exon and 3'UTR exon. **(e)** NVC (Nucleotide versus cycle) plot to check nucleotide composition bias showed A% =T%, C%=G%, which value is around 25%. **(f)** GC content distribution of reads. **(g)** Reads RPKM saturation analysis. Transcripts with expression level ranked below 25 percentile (Q1), between 25 percentile and 50 percentile (Q2), between 50 percentile and 75 percentile (Q3) and above 75 percentile (Q4) stabilized as sequencing depth increased, which measured how the RPKM deviates from real expression level. **(h)** Scatterplot of gene expression correlation in one of three representative samples by two methods: de novo analysis and genome-based analysis. Pearson's correlation in all samples r > 0.5.

Supplementary Fig.7

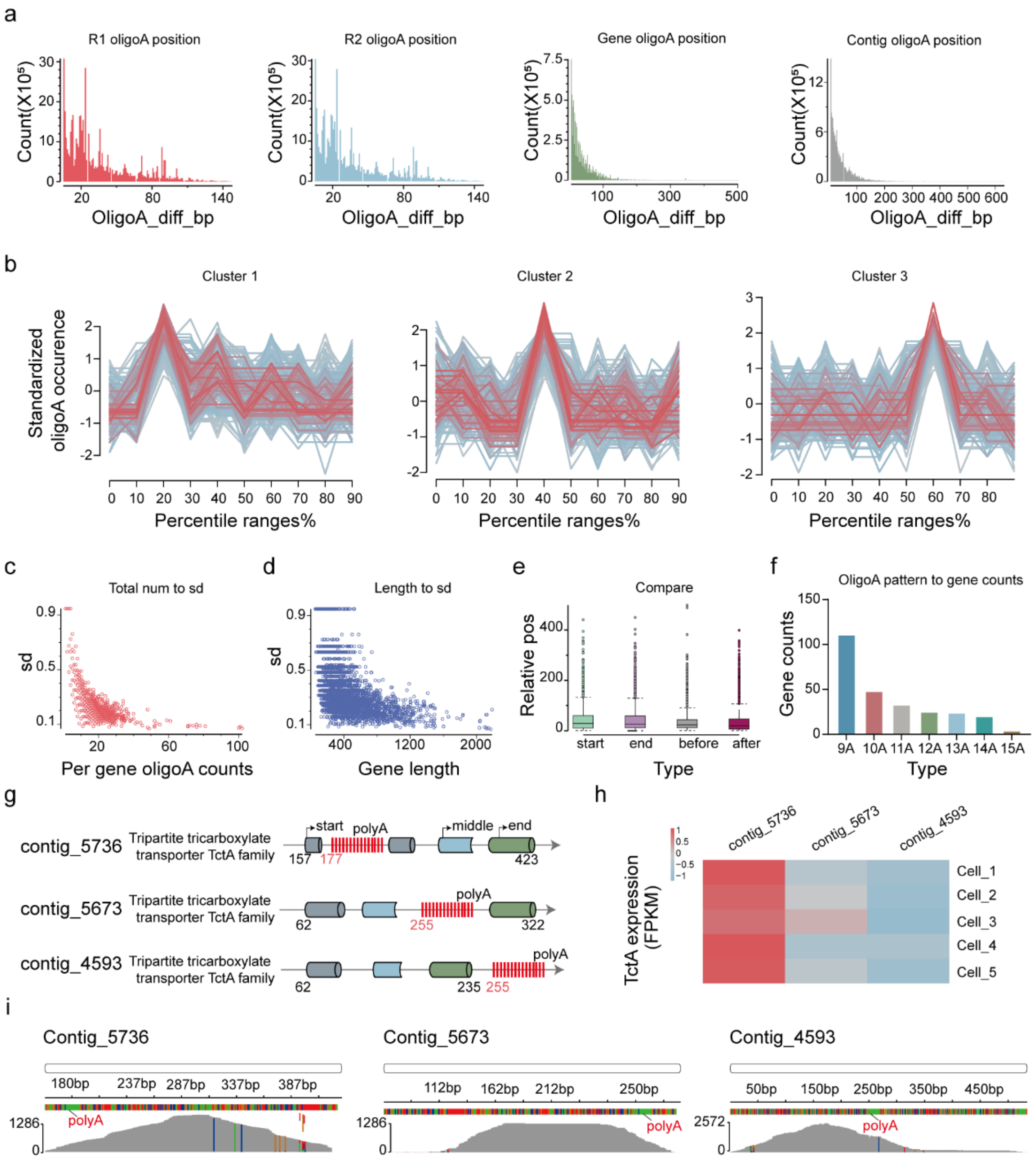


**Supplementary Fig.7 Distribution of oligoA in the *Bacillus licheniformis* transcriptome.** **(a)** Consecutive three adenylate residues (OligoA) positions were calculated in reads, genes and contigs level. **(b)** OligoA occurrence pattern within different percentile ranges of the gene length by Mfuzz analysis. Calculating proportion of OligoA count within different percentile ranges versus total OligoA count in each gene. OligoA occurrence was analysed based on proportion. Three representative OligoA occurrence pattern indicated that diverse OligoA (three adenylate residues) sites located in de novo assembly genome. **(c)** Scatterplot of standard deviation versus OligoA count in each gene. Percentages of OligoA positions within different percentile ranges was used to calculate standard deviation. **(d)** Scatterplot of standard deviation versus gene length. **(e)** Boxplot of OligoA positions within the gene, and the positions before and after the gene. 'before' is OligoA positions near to the gene start positions and 'after' is OligoA positions near to the gene end positions. **(f)** Bar plot of gene polyadenylation degree versus corresponding gene count. 9A,10A… are genes with polyadenylation level of nine, ten… adenylate residues, etc. **(g)** Illustration for oligoA detection: Polyadenylation (15 adenylate residues) in TctA gene located in three contigs from de novo assembly genome. Numbers colored black are gene start and end positions. TctA gene in each contig is split evenly into three parts according to gene total length: start part, middle part and end part. Numbers colored red indicate start positions of the polyA tracts (15 adenylate residues) according to the de novo assembly. **(h)** Gene expression heatmap (FPKM) of TctA with different polyA site in five scRNA samples treated with citric acid. **(i)** Visualization of reads coverage on TctA gene polyA sites by IGV softerware.

Supplementary Fig.8

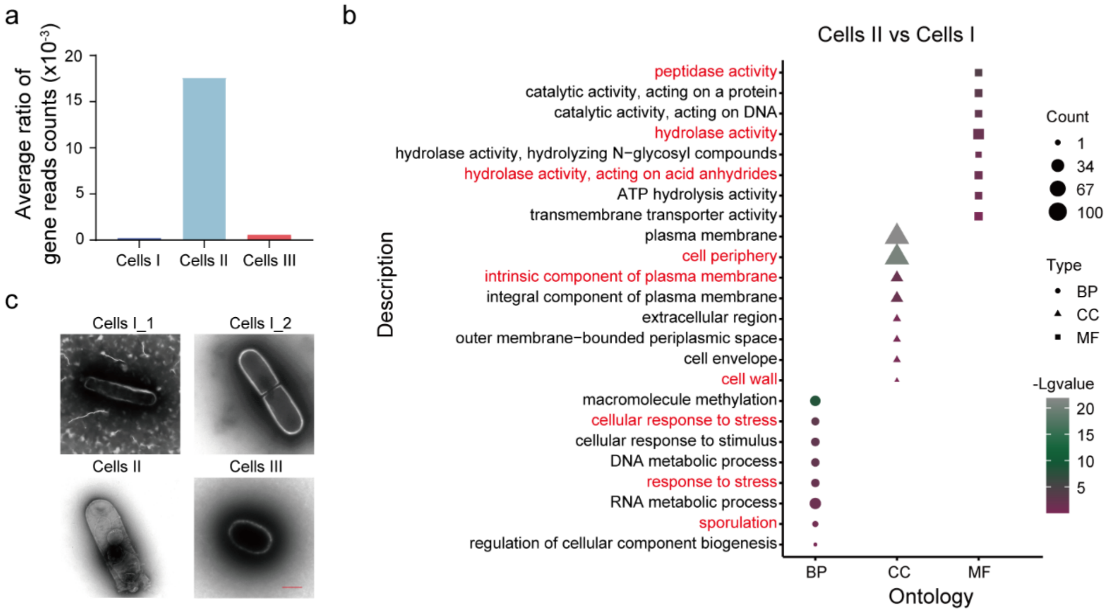


**Supplementary Fig.8 Cell status and citric acid upregulated gene pathways. (a)** Average ration of protein coding gene reads count in cells I, cells II and cells III. **(b)** Gene ontology (GO) analysis of Biological Process (BP), Molecular Function (MF), Cell Component (CC) for upregulated genes from cells II versus cells I. **(c)** Transmission electron microscope images of cells I, cells II and cells III. Scale bar, 0.5μm.

Supplementary Fig.9

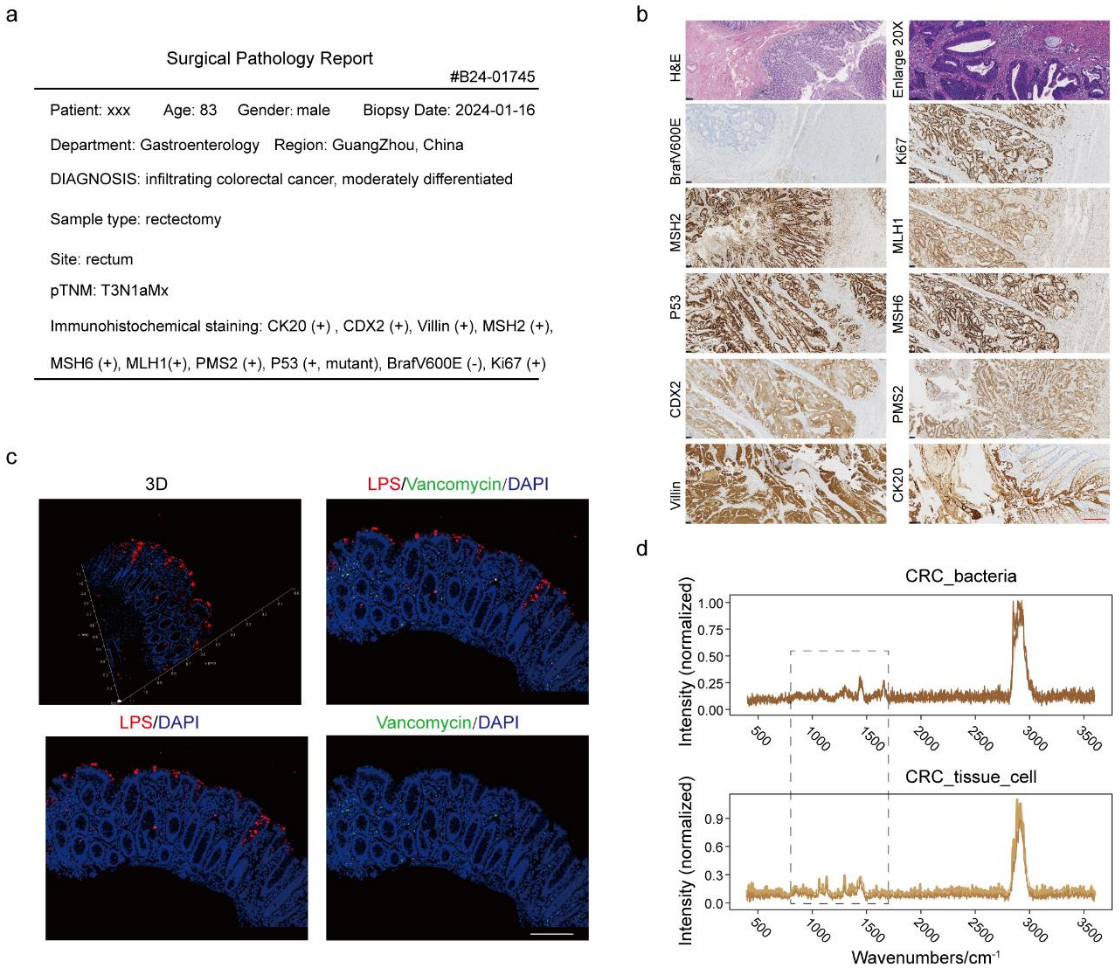


**Supplementary Fig.9 Surgical pathology report and marker gene immunostaining of tissue sections**. **(a)** Surgical pathology report to the colorectal tissue donor. The report provided key information including age, gender, diagnosis and the city where the patient lived. **(b)** Hematoxylin-eosin staining and immunohistochemical staining of tissue sections. CK20, CDX2, Villin, MSH2, MSH6, MLH1, PMS2, P53, BrafV600E, and Ki67 are colorectal cancer biomarkers. Scale bar, 200μm. **(c)** Immunofluorescence staining with antibodies against lipopolysaccharide (LPS) and vancomycin. Immunofluorescence images obtained by the laser layer scan were merged into 3D image. Scale bar, 200μm. **(d)** Raman spectra of colorectal cancer cells and microbes in tissue. The spectra are averages of three to five individual spectra and normalized to the intensity of the strongest feature in each spectrum.

Supplementary Fig.10

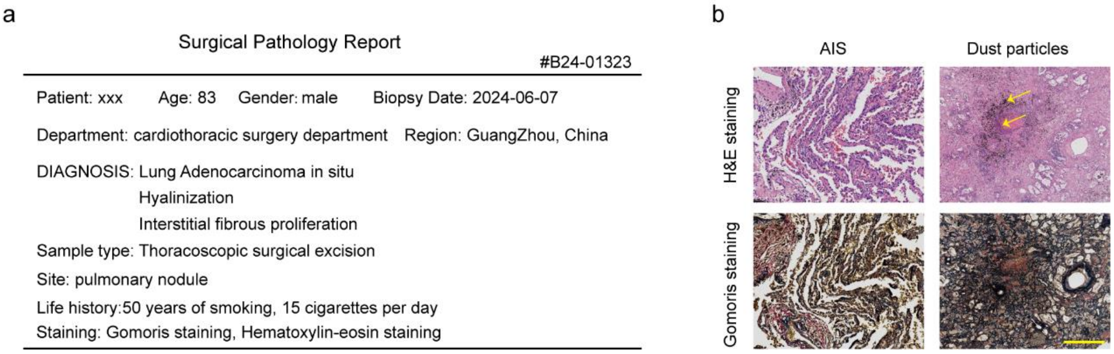


**Supplementary Fig.10 Surgical pathology report and H&E and Gomoris staining of lung tissue sections**. **(a)** Surgical pathology report to the lung tissue donor. The tissue of lung adenocarcinoma was obtained from the same patient when cancer metastasized to the lungs. The report provided key information including smoking history. **(b)** H&E and Gomoris staining of lung AIS (adenocarcinoma *in situ*) exhibited wavy arrangement of lung fibers. H&E and Gomoris staining of paracancer tissue of lung adenocarcinoma showed dust particles. Scale bar, 200μm.

**Movies S1** The procedure for running prepared samples and the software operation for automated imaging, spectral acquisition, and cell LIFT ejection.

**Movies S2** Automated single-cell LIFT ejection of TADA-labeled *Sphingomonas sp000797515* in fluorescent field.

**Movies S3** Automated LIFT ejection of single *Bacteroides xylanisolvens* suspension. After individual cells were automatically identified, these 34 cells were automatically ejected into receiver within 50 seconds.

**Movies S4** Automated single-cell LIFT ejection of salvia microbes. After automated recognition and manual selection, the selected cells were precisely ejected.

**Table S1** The obtained single-cell assembly genome of *Megamonas funiformis* was annotated by Bakta software.

**Table S2** The obtained single-cell assembly genomes from saliva were performed species annotation using GTDB-Tk and gene quality assessment using CheckM.

**Table S3** The OligoA in transcripts with three or more adenylates was calculated after transcriptome annotation.
